# HAT1 drives a gene-metabolite circuit that links nutrient metabolism to histone production

**DOI:** 10.1101/664615

**Authors:** Joshua J. Gruber, Benjamin Geller, Andrew M. Lipchik, Justin Chen, Ameen A. Salahudeen, Ashwin N. Ram, James M. Ford, Calvin J. Kuo, Michael P. Snyder

## Abstract

The energetic costs of duplicating chromatin along with DNA replication are large and therefore likely depend on nutrient sensing checkpoints and metabolic inputs. By studying chromatin modifiers regulated by epithelial growth factor, we identify histone acetyltransferase 1 (HAT1) as an induced gene that enhances cell proliferation by coordinating histone production with glucose metabolism. In addition to its canonical role as a cytoplasmic free histone H4 acetyltransferase, a HAT1-containing complex binds specifically at promoters of H4 genes. HAT1 stimulated acetate delivery and consumption at H4 promoters to drive S-phase H4 transcription. This required the presence of a histone H4-specific promoter element in the region of HAT1 chromatin binding. These data describe a feed-forward circuit whereby HAT1-dependent capture of acetyl-groups drives further H4 production to support growth-factor dependent proliferation. These findings also extend to human disease and animal models, as high HAT1 levels associate with poor outcomes across multiple cancer types.

## Introduction

There are approximately 3×10^7^ nucleosomes per human diploid nucleus (Alberts, 2002), all of which must be duplicated to capture and compact newly replicated DNA during S phase, at an estimated bioenergetic cost of ~28,000 molecules of ATP per nucleosome (Lynch and Marinov, 2015). Thus, the cellular decision to commit to cell division implies access to sufficient biosynthetic resources to meet this demand. With the exception of histone H2B (Zheng et al., 2003), little work has addressed how nutrient metabolism regulates chromatin replication.

Nucleosomes are composed of the core histone octomer (two copies each of histones H2A, H2B, H3, H4), wrapped by ~147 bases of DNA. Whereas histone variants exist for H2A, H2B and H3, only the histone H4 protein sequence remains invariant in mice and humans (Marzluff et al., 2002), indicating that H4 is a core subunit of all nucleosomes and therefore may represent a key focal point for regulation. Indeed, newly synthesized histone H4 is di-acetylated on lysines 5 and 12 by the cytosolic histone acetyltransferase 1 (HAT1) enzyme (Parthun, 2013). However, the function of this histone mark remains enigmatic, in part because it is transient, being removed 20-30 minutes after insertion into chromatin (Annunziato, 2013; Annunziato and Seale, 1983; Hammond et al., 2017; Nagarajan et al., 2013).

Recent work has integrated histone acetylation with the metabolic state of the cell. The majority of acetyl groups destined for incorporation into chromatin modifications are produced *de novo* from glucose (Wellen et al., 2009), the primary nutrient utilized by proliferating cells. Two metabolic enzymes appear to control the fate of acetyl groups destined for incorporation into chromatin. ATP citrate lyase (ACLY) generates acetyl-Co-enzyme A from citrate, a product of mitochondrial metabolism exported to the cytosol (Zaidi et al., 2012). In addition, an acetyl-Co-A salvage pathway controlled by the Acyl-CoA short chain synthetase family member 2 (ACSS2) can regenerate acetyl-Co-A from acetate at chromatin (Li et al., 2017; Mews et al., 2017), and contributes to histone acetylation in the absence of ACLY (Zhao et al., 2016) or under hypoxic conditions (Bulusu et al., 2017; Kamphorst et al., 2014; Schug et al., 2015), and is important for tumor growth (Comerford et al., 2014).

The current study is built on the observation that HAT1 is a growth factor-regulated HAT, and is required for growth factor and glucose-stimulated proliferation. This led to the identification of a nuclear role for HAT1 holoenzyme at H4 promoters, where HAT1 binding is critical for transcription of histone H4 genes. Because HAT1-induced histone acetylation is dependent on glucose availability, we conclude that HAT1 integrates nutrient availability by acetyl-group transfer to histone promoters to coordinate histone production with nutrient metabolism.

## Results

### HAT1 is an EGF-stimulated transcript required for EGF-dependent proliferation

Previous work identified EGF as the most potent stimulator of cell division in mammary cell lines (Gruber, 2018). Human mammary epithelial cells immortalized with telomerase (hTert-HME1) were dependent on epithelial growth factor (EGF) for log-phase cell division (Fig 1A). When EGF was removed from the culture media cell division continued, but at a diminished rate (Fig. 1A). To determine epigenetic regulators contributing EGF-dependent growth transcriptome analysis by RNA sequencing was performed after 2 passages (6 days) in either EGF-stimulated versus EGF-free media. Overall, there were 7117 genes differentially expressed between +/− EGF conditions (Fig. 1B, FDR 0.1), of which 4313 were up-regulated and 2804 were down-regulated in +EGF compared to –EGF conditions.

**Figure 1:**
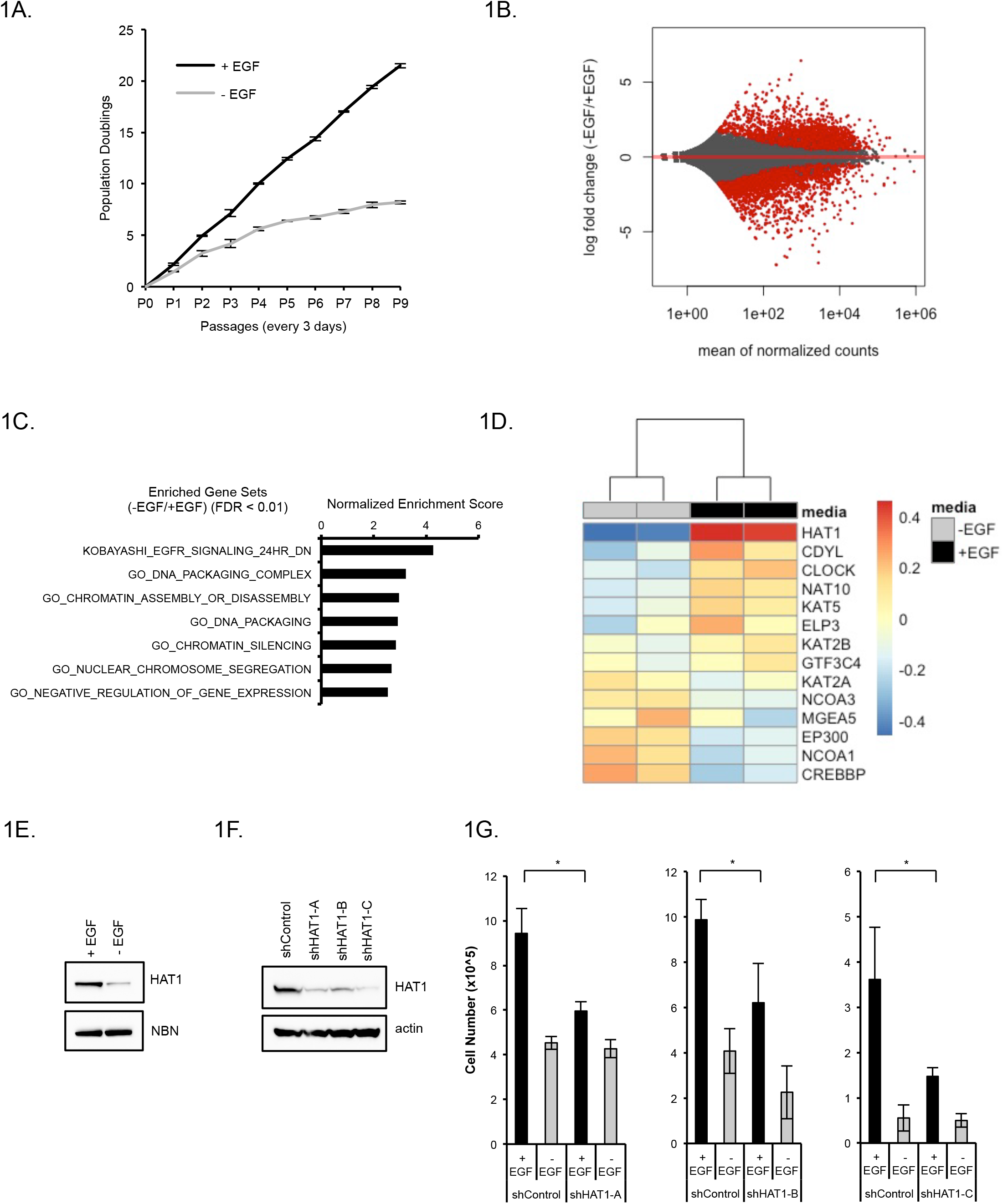
HAT1 is an EGF-regulated gene required for EGF-induced proliferation. A. hTert-HME1 cells were serially passaged in the presence or absence of EGF and population doublings +/− SD were measured. n=3 in each group. B. Gene expression profiling by RNA-sequencing of hTert-HME1 cells grown in the presence of absence of EGF for two passages. Significantly regulated genes (FDR < 0.1) are red. C. Gene set enrichment analysis was performed on differentially expressed genes identified in B. D. GO terms were used to select genes annotated to contain histone acetyltransferase activity and gene expression values are represented by heatmap. E. MCF10A cells were grown in the presence or absence of EGF for 1 passage then immunoblots were performed. F. hTert-HME1 cells were infected with lentiviruses expressing either non-targeting control shRNA or three different shRNAs targeting the HAT1 transcript. Immunoblotting was performed. G. Cells from 1F were used for proliferation assays in the presence or absence of EGF. Error bars indicated SD. * p < 0.05.

When gene set enrichment analysis was performed on the differentially expressed genes, multiple gene sets involved in chromatin biology were among the top candidates (Fig. 1C). We specifically focused on epigenetic regulators as a mechanism to understand the broad differences in gene expression observed. Gene ontology analysis identified 14 confirmed or putative genes that encoded proteins with HAT activity that were selected for further analysis. Of these genes, histone acetyltransferase 1 (HAT1) was the most strongly differentially expressed between +/− EGF conditions (log_2_ fold change 1.3; FDR < 1 x10^6^; Fig. 1D). Immunoblotting confirmed that HAT1 protein levels were also stimulated by EGF and down-regulated in EGF-free conditions in an independent but similar cell line MCF10A (Fig. 1E).

As HATs in general confer positive effects on gene expression it was tested whether HAT1 contributed to cell proliferation in EGF-stimulated conditions. Three independent shRNAs were used to reduce HAT1 protein levels in hTert-HME1 cells, compared to a control shRNA (Fig. 1F). Each shRNA targeted to HAT1 diminished proliferation in EGF-stimulated conditions, compared to control shRNA-treated cells (Fig. 1G). In contrast, no significant difference was observed between HAT1 and control shRNAs in the EGF-free condition (Fig. 1G). These data indicate that HAT1 provides a critical function in EGF-stimulated cell proliferation, but is not required for basal proliferation without EGF.

### HAT1 holoenzyme binds histone H4 promoters

We sought to understand the molecular mechanism by which HAT1 contributed to cell proliferation. HAT1 is a cytoplasmic acetyl-transferase of newly synthesized free histone H4, which it di-acetylates on lysine 5 and 12 (Parthun, 2013). It is subsequently imported into the nucleus together with the H3/H4 dimer and Rbap46/48 (Keck and Pemberton, 2012). Although HAT1 has been localized to the nucleus, its role in the nucleus remains poorly understood. To assess if HAT1 has chromatin-binding properties we performed chromatin immunoprecipitation followed by high-throughput sequencing (ChIP-seq). Compared to input control, antibodies to HAT1 enriched only 7 genomic loci (Fig. 2A, B; FDR < 0.01). Surprisingly, all of these loci mapped to a one megabase segment of chromosome 6, which is the Hist1 locus that encodes 35 replication-dependent histone genes. Examination of the location of the HAT1 ChIP-seq peaks showed that 6 of them localized within 400 bases of histone H4 gene transcription start sites (Fig. 2B, C). One localized downstream of the Hist1H2BC/H2AC locus, consistent with a potential enhancer site.

**Figure 2:**
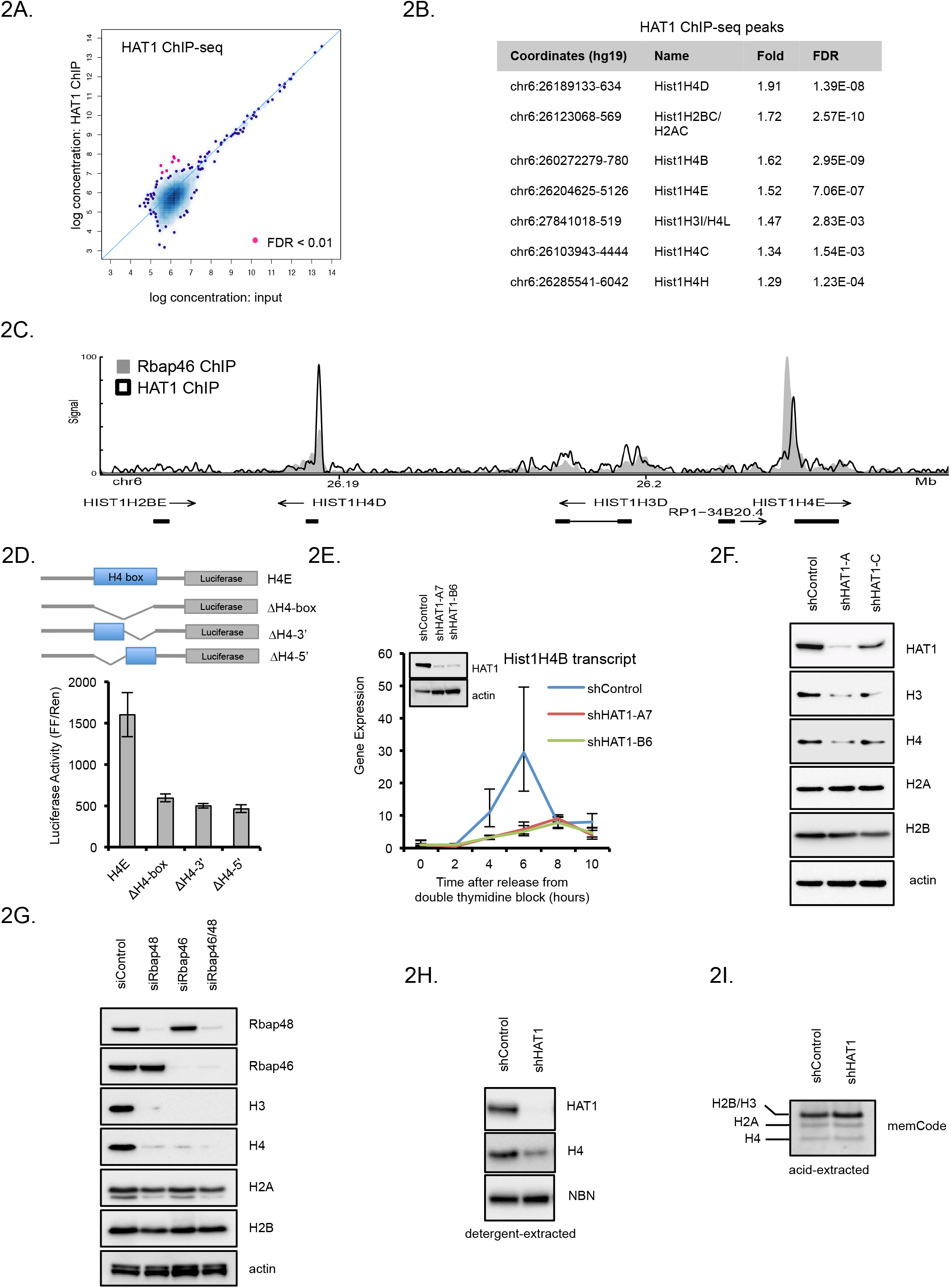
HAT1 binds are regulates transcription of histone H4 genes. A. Antibodies to HAT1 were used for ChIP-seq and significant binding was computed compared to libraries made from input material. B. Table of the 7 loci identified to significantly enrich with HAT1 antibodies compared to input control by ChIP-seq. C. Signal track of ChIP-seq data using antibodies to HAT1 or Rbap46 at the hist1 locus on human chromosome 6. D. (upper) Schematics of Firefly luciferase constructs containing 500 bp (−1 to −500) of the Hist1H4E promoter and the indicated complete or partial deletions of the H4-box. ΔΗ4-box indicates that the 17 bp H4-box sequence was deleted. ΔΗ4-3’ indicates the 3’ portion of the H4-box was deleted. ΔΗ4-5’ indicates that the 5’ portion of the H4-box was deleted. (lower) Normalized luciferase activity (Firefly/Renilla) for each luciferase construct after transfection to 293T cells. A Renilla luciferase construct driven by a TK promoter was co-transfected with the Firefly luciferase constructs to normalize for transfection efficiency. E. (inset) Immunoblot of HAT1 and actin levels in stable cell lines containing either control or two independent HAT1 shRNAs. (panel) shRNA containing cell lines were synchronized at the G1/S transition by double-thymidine block then released into S-phase and cells were harvested at the indicated time points for Hist1H4B gene expression analysis by qRT-PCR. F. hTert-HME1 cells were acutely infected with either control or two independent HAT1 shRNAs and maintained in culture for six days, then SDS-PAGE and immunoblotting was performed after detergent extraction. G. hTert-HME1 cells were transfected with the indicated siRNAs, then harvested by detergent extraction three days later and immunoblotting was performed. H. hTert-HME1 cells were infected with control or HAT1-targeting shRNAs and maintained in culture for six days, then half of the cells were extracted with detergent then SDS-PAGE and immunoblotting was performed. I. Half of the cells were treated as in 2H then histones were acid-extracted, fractionated by SDS-PAGE and stained by memCode blue.

To perform independent confirmation that the HAT1 holoenzyme associates with histone H4 promoters, we examined other proteins components of the HAT1 complex. Rbap46 is a bona fide HAT1 interactor that also binds free histone H4, likely presenting its N-terminus to HAT1 for acetylation (Murzina et al., 2008). ChIP-seq using antibodies to Rbap46 identified a much broader chromatin binding profile (30475 loci, FDR < 0.05; Fig. S1A) compared to HAT1 ChIP-seq, consistent with its known roles in other chromatin remodeling complexes including NuRD (Basta & Rauchman 2014). However, histone H4 gene promoters were among the most highly enriched loci (Fig. S1A) and Rbap46 ChIP-seq signal at the hist1 locus overlapped tightly with the HAT1 ChIP-seq signal, showing specific enrichment at the H4 promoters (Fig. 2C). Therefore, the HAT1 holoenzyme localizes specifically to histone H4 gene promoters.

DNA sequences of the H4 promoters were examined to better understand the specific chromatin-binding pattern of the HAT1 holoenzyme complex. Sequence alignments of the H4 promoters identified a highly conserved genetic element of 17 base-pairs in length that is approximately 80 bases upstream of the ATG start codons (Fig. S1B) and contained within the previously described H4 subtype specific consensus sequence (Heintz, 1991; Osley, 1991). We termed this genetic element the ‘H4 box.’ A position weight matrix using the H4 box was used to query for similar motif elements in the accessible DNA elements of the hTert-HME1 cell line derived from ATAC-seq data (Fig. S1C). This led to the identification of only H4 gene promoters (FDR < 0.01) indicating that this motif is restricted to H4 gene promoters throughout the accessible genome. Luciferase assays were used to confirm that the H4 box was critical for transactivation of the H4 promoter. Whereas the wild-type Hist1H4E promoter gave robust Firefly luciferase activity, complete or partial deletions of the H4-box robustly diminished this activity (Fig. 2D) when normalized to co-transfection of a control Renilla luciferase construct. Therefore, histone H4 promoters have a functional, unique DNA promoter element that may underlie the specific recruitment of HAT1 complexes to these loci.

### HAT1 is critical for H4 transcription in S phase

Given that the HAT1 holoenzyme complex was specifically recruited to H4 promoters, the functional significance of HAT1 directly regulating H4 gene transcription was tested. Replication-dependent histone genes, including H4, are preferentially transcribed during S-phase of the cell cycle. Therefore, control and HAT1 shRNA treated cells were synchronized at the G1/S border then released into S phase and hist1H4B RNA transcripts were measured. Depletion of HAT1 with two shRNAs caused a defect in S phase accumulation of hist1H4B transcripts compared to control shRNA (Fig. 2E). To confirm that cells depleted of HAT1 properly progressed through S-phase, cyclin E, cyclin A and phospho-H3 were immunoblotted as markers of the G1/S transition, S phase and M-phase, respectively (Fig. S1B). This confirmed that proper degradation of cyclin E, induction of cyclin A, and appearance of phospho-H3 in both control and HAT1-depleted cells after release from double-thymidine block. Immunoblots also confirmed a defect in histone H4 protein levels in cells treated with HAT1 shRNAs, compared to control (Fig. S1B). These data indicate that HAT1 is necessary to stimulate H4 gene transcription during S phase.

Given the observed effects of HAT1 on H4 transcription we also examined the levels of other core histones upon HAT1 depletion. Acute infection of hTert-HME1 cells with lentiviral shRNAs targeting HAT1 led to substantial depletion of histone H3 and H4, compared to control shRNAs, with preservation of H2A and H2B (Fig. 2F), consistent with coordinated post-translation regulation of H3/H4 dimers (Cook et al., 2011). This phenotype was also true for the HAT1 holoenzyme as siRNAs targeting Rbap46 or Rbap48 both caused decreased H3/H4 dimers without a significant change in H2A or H2B levels, compared to control siRNA (Fig. 2G). These were confirmed to be free H3/H4 dimers, rather than nucleosomal H3/H4, as determined by detergent and acid histone extraction, respectively (Fig. 2H and 2I). Thus, HAT1 promotes accumulation of free histone H3/H4 dimers without affecting levels of histones embedded in nucleosomes.

### HAT1-stimulated acetyl groups drive transcription via a H4 promoter element

We further examined the chromatin state of the hist1 locus by determining chromatin accessibility by ATAC-seq and mapping of the HAT1-dependent histone acetylation sites (H4 lysines 5 and 12) by ChIP-seq in six cell lines (3 shControl and 3 shHAT1 treated; Fig. S2A). All highly transcribed H4 genes at the hist1 locus had ATAC-seq peaks at their promoters consistent with an open chromatin state (Fig. 3A). These peaks were unchanged after HAT1 depletion (data not shown), indicating that HAT1 is not a major determinant of chromatin accessibility at H4 promoters.

**Figure 3:**
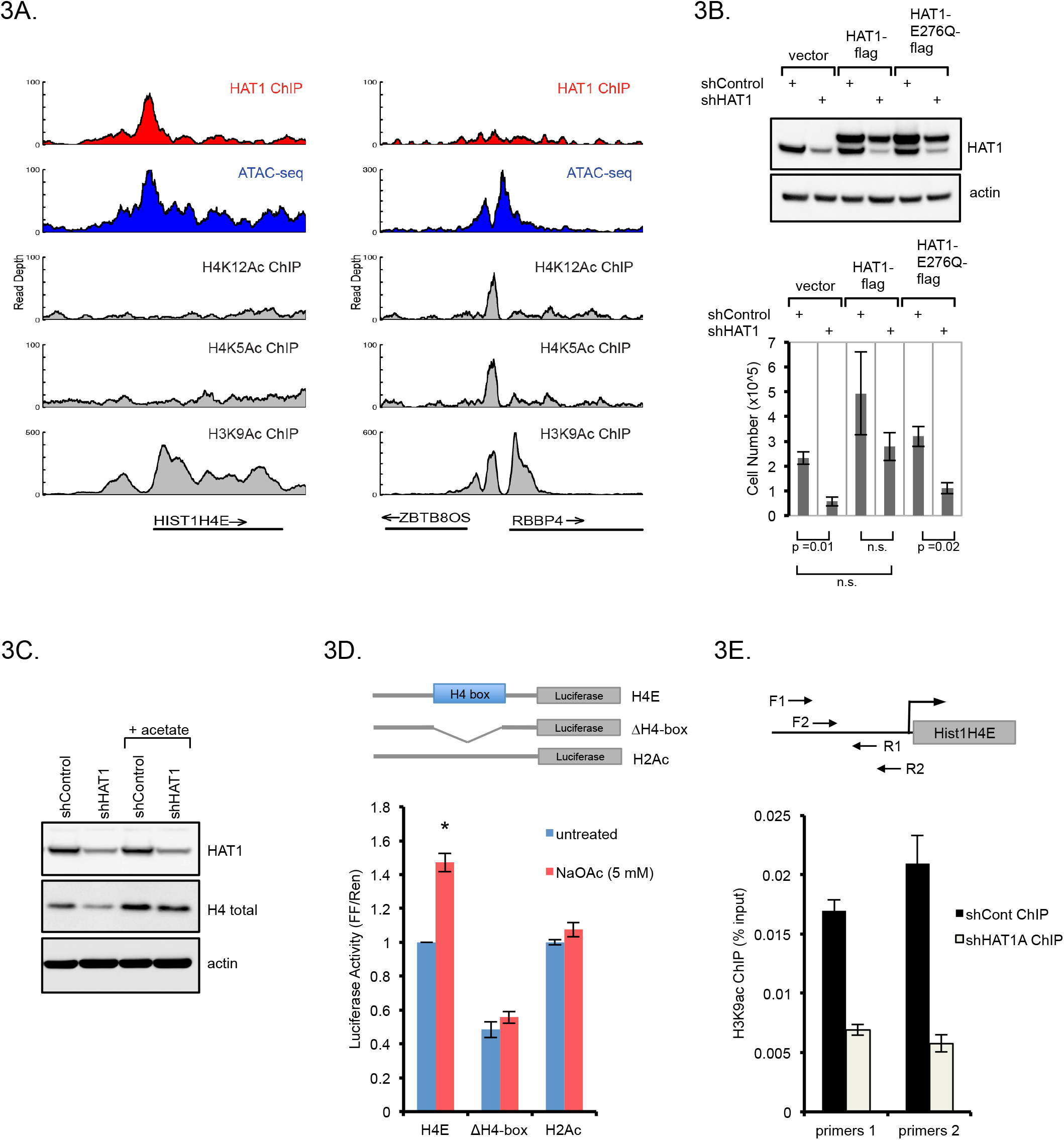
HAT1 promotes acetyl-transfer reactions to histone H3 at H4 promoters. A. Genome viewer tracks of read depth at the Hist1H4E (left) and the ZBTB8OS/RBBP4 (right) loci for HAT1 ChIP-seq, ATAC-seq, and histone modification ChIP-seq with antibodies that recognize acetylated lysines on H4K5, H4K12 and H3K9. B. hTert-HME1 cells were treated with lentivirus expressing control or HAT1 shRNAs and also with lentiviruses expressing HAT1-flag, HAT1-E276Q-flag or vector control. (upper) Protein was fractionated by SDS-PAGE and immunoblotting was performed. (lower) Proliferation was measured by cell counting +/− SD, n=3. C. hTert-HME cells were infected with control or HAT1 targeted lentiviral shRNAs and maintained in culture for 6 days. Sodium acetate (5 mM) was added on day 4. Proteins were fractionated by SDS-PAGE and immunoblotted. D. Luciferase constructs were designed as indicated with 500 bp of the Hist1H4E promoter cloned upstream of Firefly luciferase. ΔΗ4-box indicates that the 17 bp H4-box sequence was deleted. As a control 250 bp of the Hist1H2Ac promoter was used. Constructs were co-transfected with a Renilla luciferase plasmid to 293T cells and luciferase activity +/− SD was measured 24 hours later. n=3. E. hTert-HME1 cells were infected with lentiviral control or HAT1-targeted shRNAs then six days later crosslinking and ChIP-qPCR was performed with antibodies to H3K9Ac and primer sets to the Hist1H4E promoter as indicated. Error bars indicated SD, n = 3.

Because there were no changes in chromatin accessibility at the H4 promoters, we next investigated histone acetylation at these genes as a potential mechanism for HAT1 regulation. ChIP-seq for the HAT1-dependent H4K12Ac and H4K5Ac marks identified 2590 +/− 3163 and 14012 +/− 9599 (mean +/− SD; FDR < 0.01) peaks, respectively, across six cell lines (3 shControl and 3 shHAT1). However, there was no enrichment for H4K12Ac or H4K5Ac ChIP-seq signal at histone H4 promoters, despite identification of these marks at promoters elsewhere in the genome (representative examples of the Hist1H4E and ZBTB8OS/RBBP4 loci are shown in Fig. 3A). This is consistent with the high acetyltransferase activity of HAT1 towards free H3/H4 dimers, but lack of acetyltransferase activity towards nucleosomal H4, as previously published (Parthun et al., 1996). Indeed, there was no significant alteration in genome-wide H4K12Ac or H4K5Ac ChIP-seq signal between control and HAT1-depleted cell liens (Fig. S2B). Therefore, it is unlikely that HAT1-induced acetylation of nucleosomal H4 at H4 promoters affects transcription from these promoters.

As the HAT1-dependent histone modifications were not detected at the H4 promoters, we decided to test if the acetyltransferase activity was important in this system. The HAT1-depleted cell lines were rescued with cDNAs encoding either wild-type HAT1 or HAT1 with an E276Q mutation that has been shown to impair acetyltransferase function. The HAT1 cDNAs carried silent mutations to disrupt the shRNA target site to allow expression after depletion of endogenous HAT1. Whereas wild-type HAT1 rescued proliferation of HAT1-depleted cells, the HAT1-E276Q mutant did not (Fig. 3B). This confirmed that the acetyltransferase activity of HAT1 is critical to its role in proliferation.

Because the acetyltransferase function of HAT1 was required to support proliferation we tested whether exogenous acetate, the product of histone deacetylation, could rescue histone H4 levels. Cells treated with control or HAT1-targeted shRNAs were cultured in the presence or absence of 5 mM sodium acetate. Exogenous acetate supplementation rescued histone H4 protein levels after HAT1 depletion (Fig. 3C). Thus, although nucleosomal H4K5Ac and H4K12Ac are not associated with H4 promoters, free acetate is sufficient to boost H4 levels in the absence of HAT1. However, acetate supplementation was not sufficient to rescue proliferation in the absence of HAT1 (data not shown), therefore, HAT1 could contribute other essential functions for cell division beyond H4 stimulation. Nevertheless, this implies that a major function of HAT1 is to supply free acetate to H4 promoters.

Next we sought to confirm that H4 promoters were indeed acetate sensitive. The hist1H4E promoter fused upstream of firefly luciferase led to robust induction of luciferase activity when transfected to 293T cells, an effect that was further stimulated by treatment with exogenous sodium acetate (Fig. 3D). In contrast, when the H4-box was deleted from the Hist1H4E promoter (ΔΗ-box), there was diminished luciferase activity compared to the wt promoter and it was no longer responsive to acetate supplementation (Fig. 3D). As a control, the Hist1H2Ac promoter, which does not contain an H4-box, was not acetate sensitive. Therefore, histone H4 promoters are acetate sensitive and this property requires the H4-box.

Free acetate has previously been described to stimulate acetylation of lysine residues on histone H3 (Gao et al., 2016). This motivated an assessment for changes in histone H3 lysine acetylation at histone H4 promoters. ChIP-qPCR identified that depletion of HAT1 led to a loss of H3 lysine 9 acetylation at the promoter of the Hist1H4E gene (Fig. 3E). This suggests that HAT1 stimulates histone H3 lysine 9 acetylation at H4 promoters.

### Elevated HAT1 levels are associated with poor outcomes in malignancies

The above data indicate that HAT1 is a positive regulator of EGF-dependent proliferation through its ability to stimulate histone H4 expression. Furthermore, growth factor signaling pathways are common driver mutations in human malignancy. This led to the hypothesis that HAT1 expression could be an important dependency of cell proliferation in human cancers. To determine whether HAT1 expression is associated with human cancer outcomes, overall survival was stratified by HAT1 expression levels in all tumor subsets in the available TCGA data. This led to the identification of six tumor subtypes (ACC, BLCA, KICH, LGG, LIHC, LUAD) wherein high HAT1 levels (upper quartile of expression) were significantly associated with inferior survival outcomes, compared to the low HAT1 expressing tumors (lowest quartile of expression; Fig. 4A). Therefore, HAT1 expression levels could be an important predictor of patient outcomes in various human malignancies.

**Figure 4:**
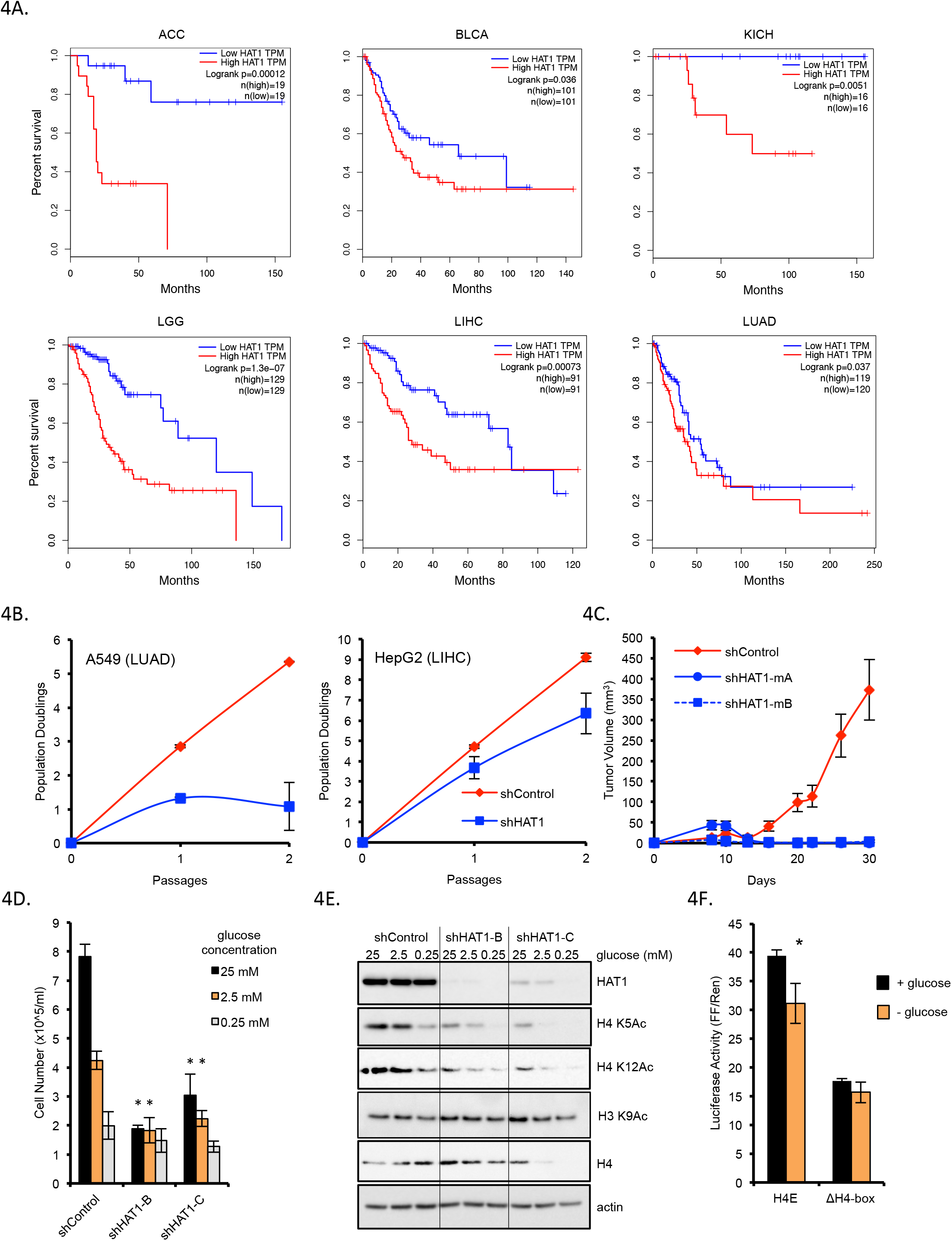
Elevated HAT1 levels are associated with poor patient outcomes in human cancer and maintain H4 expression during glucose limitation. A. Kaplan-Meier survival plots stratified by HAT1 expression levels (low = lowest quartile by z-score; high = highest quartile). Logrank p-values and total n in each group are indicated. TPM = transcripts per million (RNA-seq quantification). ACC=adrenocortical carcinoma; BLCA=bladder urothelial carcinoma; KICH=kidney chromophobe; LGG=brain lower grade glioma; LIHC=liver hepatocellular carcinoma; LUAD=lung adenocarcinoma. B. Cancer cell lines A549 (left) and HepG2 (right) were treated with control or HAT1-targeted lentiviral shRNAs and population doublings were measured. C. 4T1 mammary carcinoma cells expressing the indicated lentiviral shRNAs were orthotopically injected into bilateral mammary fat pads and tumor sizes were measured. n=5 mice in each group, each mouse bearing two tumors. Data are representative of two independent experiments. D. Stable hTert-HME1 cell lines with controls or HAT1-targeted shRNAs were grown in media containing the indicated glucose concentrations and cells were counted after three days. n=3 * p < 0.05 compared to equimolar shControl condition. E. Stable hTert-HME1 cell lines were grown in media with the indicated glucose concentrations for three days, then protein was fractionated by SDS-PAGE and immunoblotted. F. 293T cells were transfected with Firefly luciferase constructs containing either the Hist1H4E (H4E) promoter or the same promoter with the H4-box deleted (ΔΗ4-box) in glucose containing (25 mM) or glucose-free (0 mM) media. Control renilla luciferase constructs were co-transfected and luciferase activity was measured 24 hours later.

To confirm that HAT1 expression is important for human cancer cell proliferation, cell lines derived from LUAD and LIHC tumor types were treated control or HAT1 shRNAs. Depletion of HAT1 was associated with diminished proliferation in both A549 (LUAD) and HepG2 (LIHC) cancer cell lines (Fig. 4B). This suggests that HAT1 expression may be critical to support cancer cell proliferation in these cancer subtypes. In addition, HAT1 depletion impaired tumor formation *in vivo* as mice orthotopically injected with 4T1 mammary carcinoma cell lines with two independent shRNAs targeting HAT1 failed to form tumors, whereas the control shRNA-expressing 4T1 cells formed tumors robustly. Therefore, HAT1 expression engenders cancer cell proliferation and tumor formation.

Malignant cells that outgrow their local blood supply can experience nutrient limitation and metabolic stress. Glucose metabolism is a major anabolic pathway of proliferating cancer cells. To assess whether HAT1 contributed to glucose-dependent proliferation, cells were grown in varying levels of glucose-containing media. Cells with stable HAT1 depletion proliferated at a slower rate compared to control cells in both glucose-replete (25 mM) and glucose-limited (2.5 mM) conditions (Fig. 4C), whereas more profound glucose limitation equally impaired proliferation regardless of HAT1 levels. In addition, the HAT1-dependent acetylation marks on free H4 (H4K5 and H4K12) were strongly dependent on adequate glucose levels, and further reduced by HAT1 depletion (Fig. 4D). In contrast, the nucleosomal acetylation site on H3 lysine 9 was less sensitive to glucose limitation. These data indicate that HAT1 may support glucose-dependent proliferation through acetylation of free histone H4.

In contrast to the acute depletion of HAT1, which causes robust loss of histone H3/H4 dimers (Fig. 2F), stable cell lines with HAT1 depletion have normal H4 protein levels (Fig. 4D), consistent with previously published data (Nagarajan et al., 2013). However, upon glucose limitation, cells with HAT1 depletion did not maintain free H4 levels compared to control cells (Fig. 4D). To determine whether this effect occurred at the level of the histone H4 promoter luciferase assays were performed. This Hist1H4E promoter required glucose for full transcriptional activity (Fig. 4E). This effect depended on the presence of the H4-box because a Hist1H4E promoter luciferase construct with the H4-box deleted had decreased transcriptional activity and was no longer sensitive to glucose levels (Fig. 4E). Taken together, these data suggest that H4 promoters are glucose sensitive and that HAT1 is required to maintain histone H4 levels under conditions of glucose limitation.

## Discussion

Two lines of evidence support a role for HAT1 in a nutrient sensing pathway to integrate glucose metabolism with histone production. First, the HAT1-induced H4 di-acetylation marks are strongly dependent on adequate glucose levels (Fig. 4D). As glucose is the primary substrate for acetyl-Co-A generation in proliferating mammalian cells (Wellen et al., 2009) the data indicate that HAT1 captures glucose-derived acetyl groups on nascent H4. These are confirmed to be free H4 di-acetylation, rather than nucleosomal acetylation marks, because HAT1 depletion did not cause a robust change in either H4K5 or H4K12 acetylation when measured by ChIP-seq (Fig. S2B). Thus, HAT1 likely senses acetyl-Co-A availability by placing available acetyl groups on nascent H4.

A second line of evidence that supports a nutrient sensing role for HAT1 in H4 production is that H4 promoters are themselves glucose and acetate responsive. The data presented here identify a genetic element in H4 promoters that confers acetate sensitivity, which is the likely mechanism for also sensing glucose availability, because glucose is metabolized to acetyl-Co-A in proliferating cells. HAT1 contributes to this process by stimulating H3 lysine 9 acetylation at H4 promoters. As HAT1 itself does not have biochemical activity towards H3 in nucleosomes (Parthun et al., 1996), HAT1 likely stimulates H3K9 acetylation through an indirect mechanism. This could include a structural role in recruiting a functional nucleosomal H3 acetyltransferase. Alternatively, nascent H4 acetylated by HAT1 could provide a source of acetate when these histones are de-acetylated upon insertion to chromatin.

The data indicate that HAT1 drives a gene-metabolite circuit to support H4 production during growth factor stimulation. HAT1 occupies two nodal points in the circuit, as an acetyltransferase for H4 and as a chromatin-bound factor at H4 promoters. During periods of growth factor stimulation HAT1 expression is induced and the enzyme functions to capture acetyl groups by acetylating nascent H4. The HAT1/Rbap46/H3/H4 complex is subsequently imported to the nucleus via importin 4 or other karyopherin members (Campos et al., 2010). HAT1/Rbap46 is dissociated from this complex upon deposition of H3/H4 dimers at the replication fork (Agudelo Garcia et al., 2017). Thereafter, HAT1 could accumulate at H4 promoters to stimulate further production of nascent H4. As the H4 promoter requires HAT1-dependent acetyl-groups for high-output transcription, this circuit is also dependent on ongoing nutrient metabolism to support adequate acetyl-Co-A pools for H3 promoter acetylation. This feed-forward circuit likely operates to support rapid induction of H4 during S-phase to fuel chromatin replication. As HAT1 also interacts biochemically and genetically with the origin replication complex (Suter et al., 2007), this process may be directly integrated with DNA replication.

As the process of chromatin replication is essential for cell division, our data indicate that HAT1 may support this process in cancer cells. Indeed, elevated HAT1 levels were associated with poor patient survival in multiple cancer types and HAT1 was required for cancer cell proliferation in cell lines derived from these cancer types and in a mouse model of mammary carcinoma This may reflect the requirement for HAT1 to support increased H4 levels under conditions of nutrient stress, as our data suggest. Alternatively, as HAT1 deficient cells have a propensity for genome instability (Nagarajan et al., 2013; Qin and Parthun, 2002; Tscherner et al., 2012; Yang et al., 2013) it may reflect a penalty associated with ongoing DNA damage in HAT1-deficient cancer cells. Given that HAT1 depletion restricted growth of cancer cell lines, the data suggest that HAT1 could be a drug target in human cancers.

The purpose of the di-acetylation marks placed on nascent H4 has remained enigmatic since their discovery over four decades ago (Jackson et al., 1976; Ruiz-Carrillo et al., 1975). Similarly, the role of HAT1 in chromatin replication has been difficult to parse, given that nucleosome deposition can proceed without HAT1 or the HAT1-induced di-acetylation mark (Ma et al., 1998; Megee et al., 1990; Nagarajan et al., 2013; Shibahara et al., 2000; Zhang et al., 1998). Indeed, our data confirms that proliferation can proceed without HAT1, albeit at a diminished rate. Thus, di-acetylation of nascent histones may serve a metabolic purpose in providing free acetate to downstream epigenetic reactions in growth-factor stimulated cells. As a nucleo-cytoplasmic acetyl-transferase, HAT1 is ideally positioned to capture newly produced acetyl-Co-A-derived acetyl groups into nascent histones which are then shuttled to the nucleus and de-acetylated, yielding free acetate. By linking acetyl-Co-A generation to histone H4 production, HAT1 provides cells an elegant means to coordinate nutrient supply with chromatin replication in rapidly dividing cells.

## Acknowledgements

We wish to thank members of the Snyder laboratory for helpful discussions and constructive criticism of this work.

## Funding

This work used the Genome Sequencing Service Center by Stanford Center for Genomics and Personalized Medicine Sequencing Center, supported by the NIH grant award S10OD020141. J.J.G. was supported by fellowships from the Jane Coffin Childs Memorial Fund for Medical Research, Stanford Cancer Institute and Susan G. Komen Foundation, as well as funding from ASCO, the Conquer Cancer Foundation and the Breast Cancer Research Foundation. M.P.S. is supported by grants from the NIH including a Centers of Excellence in Genomic Science award (5P50HG00773502).

## Author Contributions

Conceptualization, J.J.G, B.G., J.C., A.M.L, M.P.S.; Investigation: J.J.G., B.G., A.N.R.; Validation: J.J.G., B.G., M.P.S.; Formal Analysis: J.J.G.; Resources: M.P.S.; Writing: J.J.G., J.C., J.M.F, M.P.S.; Visualization: J.J.G., B.G., J.C., A.M.L.; Supervision: J.M.F., M.P.S.; Funding Acquisition: J.J.G., M.P.S. Data and materials availability: High-throughput sequencing data has been deposited at NCBI under the GEO accession number: GSE117472.

## Declaration of Interests

M.P.S. is a founder and member of the science advisory board of Personalis, SensOmics and Qbio and a science advisory board member of Genapsys and Epinomics.

## STAR Methods

### Cell Lines

Female human cell lines MCF10A and hTert-HME1 were obtained from ATCC and were maintained in MEGM media (Lonza) in a humidified incubator at 37°C, with 5% CO2.

Glucose-free media was prepared with RPMI –glucose formulation with addition of the MEGM growth factors. HepG2 and A549 cell lines were obtained from the ENCODE consortium and maintained in MEM and F-12K medias, respectively, supplemented with 10% FBS and 1% penicillin:streptomycin.

### Transfection, cell assays

siRNA transfection was performed with 30 microliters of Lipofectamine RNAiMAX (Invitrogen), 3 mL OptiMEM media, 18 picomoles of siRNA and 5E5 – 1E6 cells in 10 cm plates. Cell proliferation was measured with a Biorad TC 10 with trypan blue. Population doublings were calculated using the formula: PD = 3.32 (log(counted) – log(plated)), where PD = population doublings, counted = number of cells counted, plated = number of cells plated. For luciferase assays, 500 bp of promoter sequence was cloned into the pGL4.23 vector (Promega) and constructs were transfected to 293T cells with the Xtremegene 9 reagent (Roche). One tenth mass equivalent of pRL-TK plasmid was co-transfected. Luciferase activity was measured with Dual-Luciferase Reporter System (cat# E1910; Promega) 24 hours after transfection. Double-thymidine block was performed by addition of 2 mM thymidine for 18 hours, followed by 9 hour wash-out with fresh media, then a second block with thymidine (2 mM) for 15 hours, then wash-out and release in the G1/S.

### RNAi and cDNAs

HAT1 shRNAs were purchased from OriGene with the following sequences:

A. GATGGCACTACTTTCTAGTATTTGAGAAG;
B. AAGGATGGAGCTACGCTCTTTGCGACCGT;
C. TCCTACAGTTCTTGATATTACAGCGGAAG.

The HAT1 cDNA was purchased from OriGene (catalog# RC209571L1) and the 5’ end was extended by a Gibson assembly reaction to add the additional 255 nucleotides to obtain a full-length clone. Mutagenesis of this cDNA was performed with the QuikChange II Site-Directed Mutagenesis kit with the following primers to make the D276Q acetylation-dead form:

CAGTTCTTGATATTACAGCGCAAGATCCATCCAAAAGCTAT, ATAGCTTTTGGATGGATCTTGCGCTGTAATATCAAGAACTG.

To mutate the shRNA-C target site mutagenesis was performed with the following primers: GGTCGCAAAGAGCGTAGCTCCGTCTTTGTTGTATTTCTCAAATACTAGAAAGTAGTGCCATCTTTCATC, GATGAAAGATGGCACTACTTTCTAGTATTTGAGAAATACAACAAAGACGGAGCTACGCTCTTTGCGACC.

### Mouse tumor studies

The 4T1 mammary carcinoma cell line was infected with lentiviral control or HAT1-targeted shRNAs (mA-GAAGCTACAGACTGGATATTA; mB-GAAGATCTTGCTGTACTATAT) in the pLKO-puro-IPTG-3xLacO vector (Sigma-Aldrich) then selected in puromycin and single cell clones were obtained by limiting dilution. To minimize clonal variation, two independent cell line clones from each shRNA were mixed in 1:1 ratio prior to injection of 50,000 cells to bilateral mammary fat pads of 4-6 week-old female Balb/c mice. At day 1 after cell injection 20 mM IPTG was added to the water bottles of all cages to induce shRNA expression and water was changed once a week with repeat addition of IPTG. Tumor size was measured by calipers.

### Immunoblots, qRT-PCR

Protein extracts were made in RIPA buffer and quantitated by BCA assay and diluted to equal concentrations. Polyacrylamide gel electrophoresis was performed on NuPAGE Novex gradient gels (Thermo Fisher) followed by wet transfer to nitrocellulose membranes. Blocking was briefly performed with 5% non-fat milk and primary antibody was incubated overnight at 4°C, then with HRP-conjugated secondary antibody (Cell Signaling) at room temperature for 1 hour followed by washing, then developed with ECL pico or femto (Thermo Fisher). For gene expression assays total RNA was isolated with All-prep (Qiagen), Dnase treated, then reverse-transcribed with Superscript III (Invitrogen). qPCR was performed with 2x KAPA SYBR Fast Master Mix on a QuantStudio Flex 6 (Applied Biosystems).

### ChIP-seq

Log-phase growth cells were crosslinked in 1% formaldehyde at a density of 5×105 cells per milliliter for 10 minutes at room temperature, then quenched with 125 mM glycine for 5 minutes, washed in PBS and snap frozen. Cells were then thawed and nuclei isolated by triton-X permeabilization followed by washing in a low detergent buffer. Then RIPA was added and sonication was performed by Branson (3 pulses of 30 seconds each with power output 4 W), followed by 14 cycles of sonication on the Bioruptor Pico. Chromatin extracts were then cleared by centrifugation and immunoprecipitation was performed with antibodies and protein A/G agarose beads overnight. The next day the beads were extensively washed with RIPA, then PBS, then resuspended in TE with 1% SDS and crosslinks were reversed overnight at 65 degrees Celsius. The next day RNase A treatment and proteinase K treatment were performed, followed by recovery of DNA with the Qiaquick spin columns. High throughput sequencing libraries were then constructed by A tailing, adapter ligation and 10-15 cycles of PCR followed by library purification, removal of PCR primer-dimers and high-throughput sequencing by HiSeq 4000. Antibodies for ChIP-seq: Rbap46 (V415) from Cell Signaling (cat# 6882); HAT1 from Abcam (cat# ab194296); Histone H4 acetyl K12 from Abcam (cat# ab46983); acetyl-Histone H4 (Lys5) from Millipore (ca# 07-327).

### ChIP-seq and ATAC-seq peak calling and analysis

Paired-end 100 bp reads were trimmed with cutadapt version 1.8.1 with flags –u −50 –U −50 –a CTGTCTCTTATACACATCTCCGAGCCCACGAGAC -A CTGTCTCTTATACACATCTGACGCTGCCGACGA -O 5 -m 30 -q 15. Bowtie (Langmead et al., 2009) version 1.1.1 was used to align trimmed reads to hg19 with flags -q --phred33-quals -X 2000 -m 1 --fr -p 8 -S --chunkmbs 400, followed by samtools sort command. Duplicates were marked with Picard-tools (Picard-tools) version 1.92, then samtools view with flags –b –f 1 -F12 –L were used to filter mitochondrial mapping reads with a bed file containing all chromosomes except chrM. SPP/phantom (Kharchenko et al., 2008; Landt et al., 2012) was run to obtain the fragment length with maximum strand cross-correlation. MACS2 (Zhang et al., 2008) callpeak function was then performed with flags -q 0.05 --nomodel –extsize=1/2 fragment length obtained from SPP. The program align2rawsignal (Consortium, 2012) (Kundaje) was used to create genome-wide signal coverage tracks with normalization to account for depth of sequencing and read mappability with flags kernel (k)=epanechnikov, fragment length (l)=150 (ATAC-seq) or (l)=1/2 fragment length from SPP for ChIP-seq, smoothing window (w)=150, normFlag(n)=5, mapFilter (f)=0. Further peak calling was performed with DiffBind (Ross-Innes et al., 2012; Stark, 2011) with summits=250 paramter contrasting ChIP-antibody to input control to obtain final enriched peak counts.

### TCGA gene expression analysis

Exploratory analysis and generation of survival plots and statistics was performed with the GEPIA resource (Tang et al., 2017).

