## Supplemental_material for "HAT1 drives a gene-metabolite circuit that links nutrient metabolism to histone production"

### Figure S1, Related to Figure 2.

S1A.

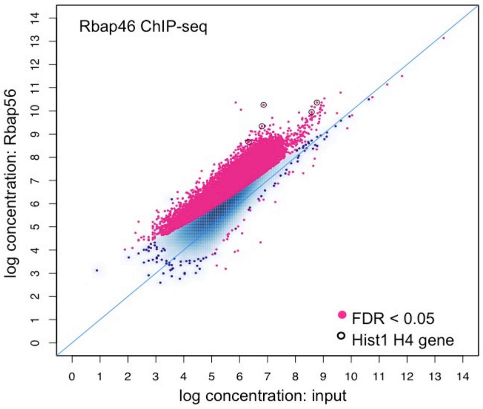


S1C.

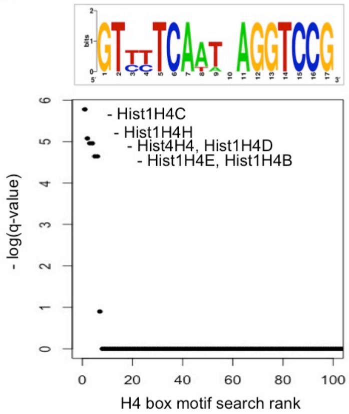


S1B.

```
Hist1H4c    509 AAAACAATGAACCAT-AAT---GAAACAGTTTTTCTTGCTCCACCCACTTGGT--GACCA
Hist1H4d    417 GCCTCAGGCCCCTCTTGGTCTCCTCC----AGTCTGAGCAAGGAGAATTGCACAGAAGG
Hist1H4B    425 CTCCCTCGCAGTCCTTAG----GTCACTGCCCCTCGAGGGGCGGAGCAAAAAGTGAGGCA
Hist1H4e    510 GGGAACTGGAGTCGTTGGTGACGTCATCCCAGTCTGATCTGTGAAGGGTAGGGCCAGCAG
Hist1H4h    556 AAAGGCTGGAGTCGTTGGTGACGTCATCCAGGGCTGGGCTATGCGGGGTGGGGCCAAGAA
consensus   601 a      g aa c Ttagt   gtca          gagc a g agaat gga  ga  a

Hist1H4c    563 AATTTGAAAAAAAAAAAAAACCGCGCCAACTCATGTTGTTTTCAATCAGGTCCGCCAAGT
Hist1H4d    473 GCGGGAACTGACTGTGGTTCCCGCCCCTTCTCATGAGGTTTTCAAACAGGTCCGTCATGC
Hist1H4B    481 GC-----AACGCCTCCTTATCCTCGCTCCCGCTTTCAGTTCTCAATAAGGTCCGATGTTC
Hist1H4e    570 GC-----AGCACCAAAGTTCCCGTATGCGCGTTTTCAGTCTTCATTTAGGTCCGAATTCC
Hist1H4h    616 GC-----AGGACCAAAGTTCCGGTATGCGCGCTTTCAGTCTTCAATTAGGTCCGAATTCC
consensus   661 gc     aa ac a aat  Ccg g    C c T  gGT tTCAat AGGTCCG    t c

Hist1H4c    623 TTGTA--------------------TTTAAGGAACTGTTTCAGTTCATACCTTCCACTGCG
Hist1H4d    533 TATTATAAAAGTGCTGC-GTGGCTCTGCCAGACGTATTGCGTTACAGATTTAACAGCTGT
Hist1H4B    536 GTGTATAAATGCTCGTG----GCTTGCTTTCTTTTCGCGTACCTGGTTTTTGTTGTCAGC
Hist1H4e    625 CGGCATATAAGAATACTACCGTCGCTTGTTTTTCAGATTTTTGCGGCTATTTTCGTTGGT
Hist1H4h    671 CGGCATATAAGGGCGTTGCTTTGCCTTGGCGTTTAGGTTTCTTAAGTTGGTTTTAGAAGT
consensus   721   g Ata a g                t          g t        a    T   a  gg

Hist1H4c    664 ATAGGAATCATGTCTGGTCGCGGCAAAGGCGGAAAAGGCTTGGGGAAGGGTGGTGCTAAG
Hist1H4d    592 GGTTTCAAGATGTCTGGCCGCGGTAAGGGCGGAAAGGGTCTAGGTAAGGGTGGCGCCAAG
Hist1H4B    592 TGGTTAGACATGTCTGGTCGCGGCAAAGGCGGTAAAGGTTTGGGTAAGGGAGGTGCCAAG
Hist1H4e    685 GTGTTGGTCATGTCTGGTCGCGGCAAAGGCGGAAAGGGACTGGGTAAAGGAGGCGCTAAG
Hist1H4h    731 TGCTTAGTCATGTCTGGCCGTGGTAAAGGTGGAAAAGGTTTGGGTAAGGGAGGAGCTAAG
consensus   781    ttaa cATGTCTGG CGcGG AAaGGcGGaAAaGG  TgGGtAAgGG GG GC AAG
```

S1D.

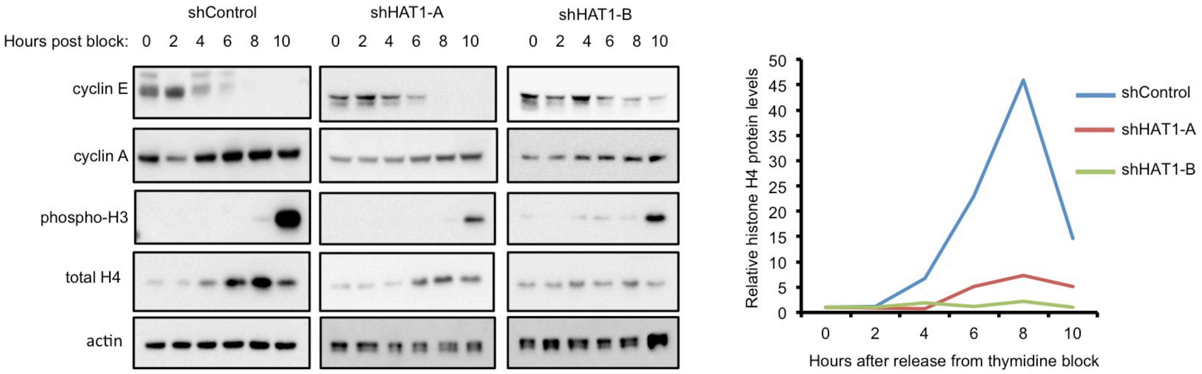

**Figure S1, Related to Figure 2**

A. Antibodies to Rbap46 were used for ChIP-seq and differentially bound loci were computed compared to input control. Loci significantly enriched by antibodies to Rbap46 are in red (FDR $<0.05$) and loci linked to Hist1 H4 genes are circled.

B. Multiple sequence alignment of Hist1H4 genes bound by HAT1 as determined by ChIP-seq. The H4-box is indicated with red underline. The H4 gene start codon is underlined in blue.

C. (upper) Position-weight-matrix of the H4-box identified in the promoters of HAT1-bound H4 genes. (lower) The H4-box position-weight-matrix was used to query the accessible chromatin landscape of hTert-HME1 cells (ATAC-seq intervals) and matching loci are ranked by significance.

D. hTert-HME1 cells containing either control or two independent HAT1 shRNAs were synchronized at the G1/S boundary by double-thymidine block then released into S-phase and collected at the indicated times. (left) SDS-PAGE and immunoblotting was performed. (right) Densitometry quantitation of H4 protein levels from the shown immunoblots.

**Figure S2, Related to Figure 3.**

S2A.

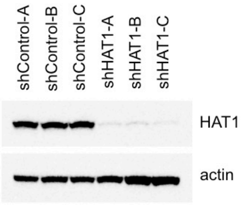


S2B.

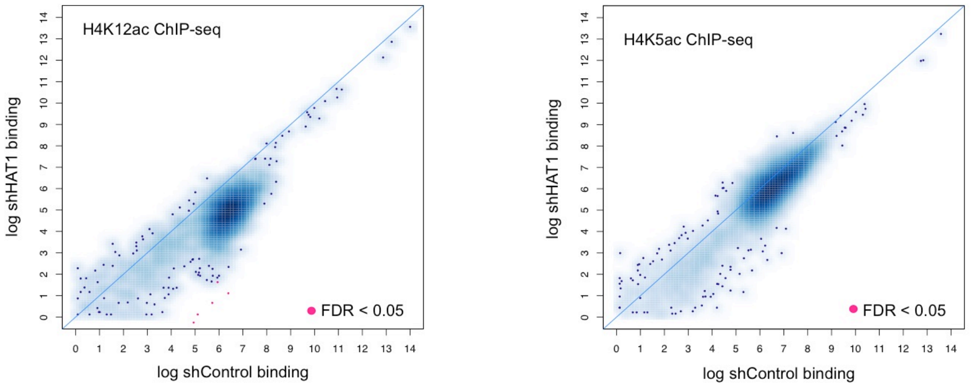


**Figure S2, Related to Figure 3.**

A. Stable cell lines expressing control shRNAs or three independent shRNAs to HAT1 were fractionated by SDS-PAGE, then immunoblotted.

B. ChIP-seq was performed with antibodies recognizing H4K12ac or H4K5ac. Enriched loci were defined compared to input control sequencing. Loci that were significantly enriched or depleted are in red (FDR <0.05).
